# Augmentation of DNA exonuclease TREX1 in macrophages as a therapy for cardiac ischemic injury

**DOI:** 10.1101/2024.02.20.581294

**Authors:** Ahmed Gamal-Eldin Ibrahim, Alessandra Ciullo, Kazutaka Miyamoto, Ke Liao, Xaviar M. Jones, Shukuro Yamaguchi, Chang Li, Alice Rannou, Asma Nawaz, Ashley Morris, Kara Tsi, Cristina H. Marbán, Jamie Lee, Nancy Manriquez, Yeojin Hong, Arati Naveen Kumar, James F. Dawkins, Russell G. Rogers, Eduardo Marbán

## Abstract

Noncoding RNAs (ncRNAs) are increasingly recognized as bioactive. Here we report the development of TY1, a synthetic ncRNA bioinspired by a naturally-occurring human small Y RNA with immunomodulatory properties. TY1 upregulates TREX1, an exonuclease that rapidly degrades cytosolic DNA. In preclinical models of myocardial infarction (MI) induced by ischemia/reperfusion, TY1 reduced scar size. The cardioprotective effect of TY1 was abrogated by prior depletion of macrophages and mimicked by adoptive transfer of macrophages exposed either to TY1 or TREX1. Inhibition of TREX1 in macrophages blocked TY1 cardioprotection. Consistent with a central role for TREX1, TY1 attenuated DNA damage in the post-MI heart. This novel mechanism—pharmacologic upregulation of TREX1 in macrophages—establishes TY1 as the prototype for a new class of ncRNA drugs with disease-modifying bioactivity.

**One Sentence Summary:** Upregulation of three prime exonuclease, TREX1, in macrophages enhances tissue repair post myocardial infarction.

## Introduction

During and after myocardial infarction (MI), the heart undergoes oxidative stress, culminating in sterile inflammation and tissue necrosis(*1*). Long-term sequelae include fibrosis, adverse remodeling, and ventricular dysfunction, leading to heart failure(*2*). At a cellular level, oxidative stress drives DNA damage through the induction of double-stranded DNA breaks, replication fork collapse, and histone displacement(*3, 4*). Although reactive oxygen species can independently trigger endoplasmic reticulum (ER) stress through the accumulation of misfolded proteins, DNA damage also contributes directly to ER stress by mediating ER-mitochondria contacts and promoting p53-induced apoptosis(*5*). DNA damage and ER stress recruit powerful innate immunity defense programs(*3, 6*). The three-prime DNA exonuclease, TREX1, serves as a vital gatekeeper for innate immunity: TREX1 senses and degrades damaged DNA, but, when its capacity to do so is exceeded, activation of the cGAS-STING pathway leads to rampant inflammation, cell stress, and senescence(*6*). The proinflammatory impact of DNA damage is particularly evident in macrophages, which orchestrate acute and chronic inflammatory responses(*7, 8*). TREX1 is induced in macrophages in response to proinflammatory stimuli; indeed, TREX1 deficiency induces an exaggerated proinflammatory response in macrophages (*9*), systemic autoimmunity(*10, 11*) and accelerated senescence(*12*). Nevertheless, the converse conjecture remains unexplored: might TREX1 augmentation be helpful in disorders of inflammation? Here, we leverage a serendipitous discovery—a novel noncoding (nc) RNA drug that upregulates TREX1—to probe the therapeutic implications of the DNA damage response in acute MI(*13*). Augmenting TREX1 in macrophages suffices to reduce infarct size in both rats and pigs. This novel cardioprotective mechanism—targeting the DNA damage response—has broad-reaching therapeutic implications for inflammatory disorders.

## Results

### Structure-activity relationship studies yield lead compound TY1

Extracellular vesicles (EVs) from cardiosphere-derived cells (CDCs) are immunomodulatory and cardioprotective(*14-20*). In probing previously-unknown bioactive constituents of CDC-EV cargo, we identified EV-YF1, a Y RNA that attenuated tissue damage in models of MI and/or hypertrophy(*21-23*). As a therapeutic candidate, however, EV-YF1 defies the structural conventions for FDA-approved ncRNA drugs (too long [56 nt] and chemically-unmodified)(*24*). We thus embarked on structure-activity optimization. **Figure 1A** shows the aligned sequences of EV-YF1 (top) and NT4, a truncated species (24 nt, bottom) which is also plentiful in CDC-EVs(*25*). Both EV-YF1 and NT4 increase the expression of the anti-inflammatory cytokine Il10 in rat bone marrow-derived macrophages (BMDM; **Fig. 1B**), an *in vitro* finding which correlates well with *in vivo* disease-modifying bioactivity(*21-23*). Using NT4 as the template, we systematically synthesized various modified versions and screened them *in vitro* for Il10 upregulation in BMDM(*21*). We first mutated the natural sequence, then included locked nucleic acids (LNAs), and finally added methylated nucleotides or adenines (**Fig. 1C**). The overall goal was to preserve bioactivity while optimizing structure, following what has been learned about structure-activity relations from successful ncRNA drug development studies(*24, 26, 27*). Substituting the first NT4 residue (guanine) with cytosine (NT4^1G-C^) yielded greater Il10 upregulation than uracil or adenine substitutions, which were largely inert (**Fig. 1D**). We thus proceeded with NT4^1G-C^, adding backbone LNAs that increase the stability of stem-loop structures(*26*). The inclusion of three alternating LNA residues on both ends of the oligonucleotide (NT4^1G-C^ 3 alt.) enhanced the functionality of NT4^1G-C^, while using all LNA nucleotides (NT4^1G-C^ 3 All) or three LNA consecutive residues on each end (NT4^1G-C^ 3 cons) abrogated bioactivity entirely (**Fig. 1E**). The optimal lead compound, NT4^1G-C^ 3 alt., was dubbed *T*herapeutic *Y* RNA *1*, or *TY1* (**Fig. 1E**). We additionally examined further modifications designed to minimize RNA degradation. Exonucleases that target the 3’ end are inhibited by adding a methyl group on the ribose sugar of the dNTP (2’-O-methylation)(*28*), or adenine residues (1 or 2 adenine additions have been shown to provide maximal stability(*29*)). In mouse BMDM, addition of a single adenine (TY1^1m^) or one methylated residue (at the 3’ terminal guanine; TY1^1A^) modestly enhanced Il10 expression (**Fig. 1F**), but this observation did not hold up in human macrophages (**Fig. 1G**). Based on these collective results, we chose TY1 for further development. To recap, relative to NT4, TY1 contains a point mutation (G to C) at its 5’end, as well as six LNA modifications at the residues underlined (**Fig. 1H**). As a control, we created a scrambled version (Scr) containing the same nucleotides as TY1 but in random order (and verified to lack sequence homology to either the mouse or human genomes), with 6 LNAs at the same positions (**Fig. 1H**). Finally, to quantify stability, we exposed TY1 and its natural precursor molecules (EV-YF1 and NT4) to ribonuclease R (RNAse R), the enzyme primarily responsible for degradation of small RNAs *in vivo*(*30-32*). TY1 was quite resistant to RNase R, unlike the natural products which were degraded by RNase R in a concentration-dependent manner (**Fig. 1I**). Finally, to assay therapeutic effects beyond Il10 production, we exposed lipoprotein S (LPS)-challenged macrophages to TY1, Scr, or vehicle. Mimicking the bioactivity of EV-YF1(*21*) and NT4(*33*), TY1 reduced the expression of stress and inflammatory markers including P21 (**Fig. 1J**), NFkb (**Fig. 1K**), and Il6 (**Fig. 1L**) in BMDM. Thus, TY1 is a shorter, more stable new chemical entity that retains the bioactivity of EV-YF1.

**Figure 1:**
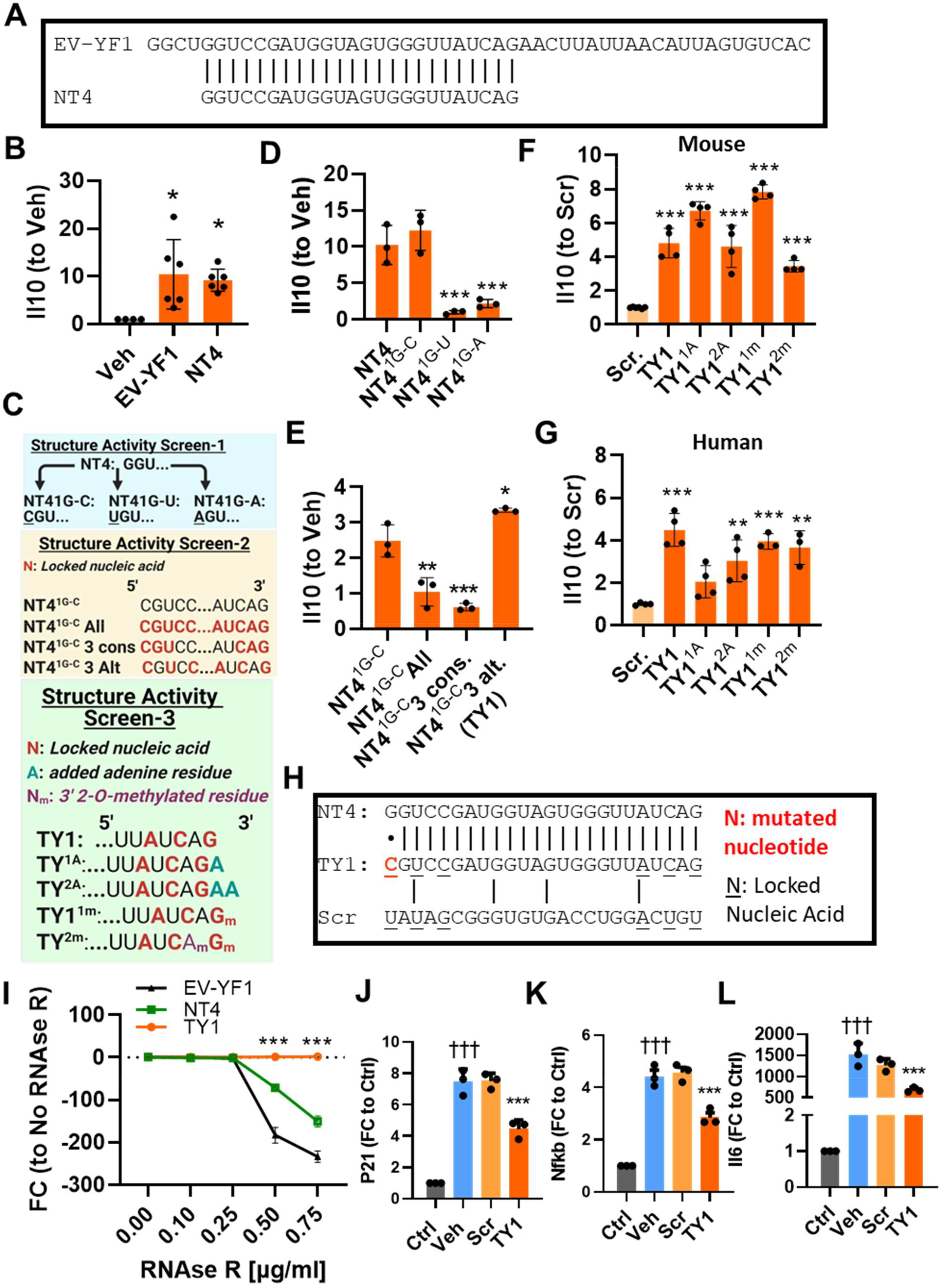
TY1 is an engineered small RNA with anti-inflammatory properties. (**A**) Alignment of EV-YF1 and NT4, both of which are abundant in the extracellular vesicles (EVs) secreted by cardiosphere-derived cells (CDCs). (**B**) As has been shown for EV-YF1, NT4 upregulates Il10 expression in rat bone marrow-derived macrophages (BMDM) compared to vehicle controls (n=6 biological replicates per group). (**C**) Schematic of three-step screening of NT4-derived structural variants for optimization of activity. Step 1 (blue shading): mutagenesis; step 2 (yellow shading), the inclusion of locked nucleic acids (LNA; phase 2); step 3, poly-adenylation and methylation of 3’ residues. (**D**-**G**), Screening of compounds was performed in murine and human macrophage using Il10 expression assays n=3-4 biological replicates per group. (**D**) Step 1 screening of different nucleotide substitutions of the most 5’ nucleotide (a guanine; all groups compared to NT4). (**E**) Step 2 screening of variants including LNAs (all groups compared to NT4^1G-C^). (**F**,**G**) Step 3 screening evaluating the adenylation and methylation at the 3’ end of TY1 in mouse, (**F**), and human macrophages,(**G**; all groups compared to TY1). (**H**), comparison of naturally-occurring NT4 and its optimized, bioinspired derivative (TY1). Also depicted is the sequence of a scrambled variant (Scr) with the same nucleotide content and LNAs, which serves as a control in panels (**F** and **G**). (**I**) Stability of TY1, NT4, and EV-YF1 exposed to increasing concentrations of the exonuclease RNAse R (n=3 biological replicates per group). BMDM pre-exposed to TY1 before lipopolysaccharide (LPS) exposure showed reduced markers of senescence (P21; **J**), and inflammation (Nfkb and IL6; **K, L**) compared to vehicle or scrambled sequence (n=3 biological replicates). Bars and line dots represent group mean and error bars represent s.d. Significance was determined by one-way ANOVA; *P<0.05; **P<0.01; ***; P<0.001. *; denotes comparison between groups and vehicle, †; denotes comparison between groups and control.

### TY1 activates genotoxic response genes and attenuates cell stress

TY1, admixed with DharmaFECT® to create lipid nanoparticles, induced profound transcriptomic changes in BMDM (heat map, **Fig. 2A**). Ingenuity Pathway Analysis implicates the DNA damage response as strongly activated by TY1 (**Fig. 2B**). Interrogating differentially-expressed DNA damage response genes identified the TREX1-cGAS/STING pathway as especially enriched (circumscribed in red; **Fig. 2C, D**). Downstream of this genotoxic stress pathway, we observed upregulation of anti-inflammatory mediators (**Fig. 2E**), the unfolded protein response (UPR) sensors of the endoplasmic reticulum (ER; **Fig. 1F**), and effector E1-E3 ubiquitin system (**fig. S1A**). Attenuation of genotoxic stress, coupled with enhanced ability to clear misfolded proteins, would logically suppress cell stress signaling more broadly, a prediction borne out by the finding that MAPK signaling was downregulated by TY1 (**fig. S1B**). DNA damage caused by oxidative injury promotes ER stress(*34*), which in turn, leads to lower protein-folding capacity and the accumulation of stress granules containing misfolded proteins(*35, 36*). Consistent with suppression of the UPR, TY1 decreased ER-stress-induced misfolded protein (aggresome) accumulation in LPS-exposed BMDM, compared to Scr or vehicle (**Fig. 2G, H**).

**Figure 2:**
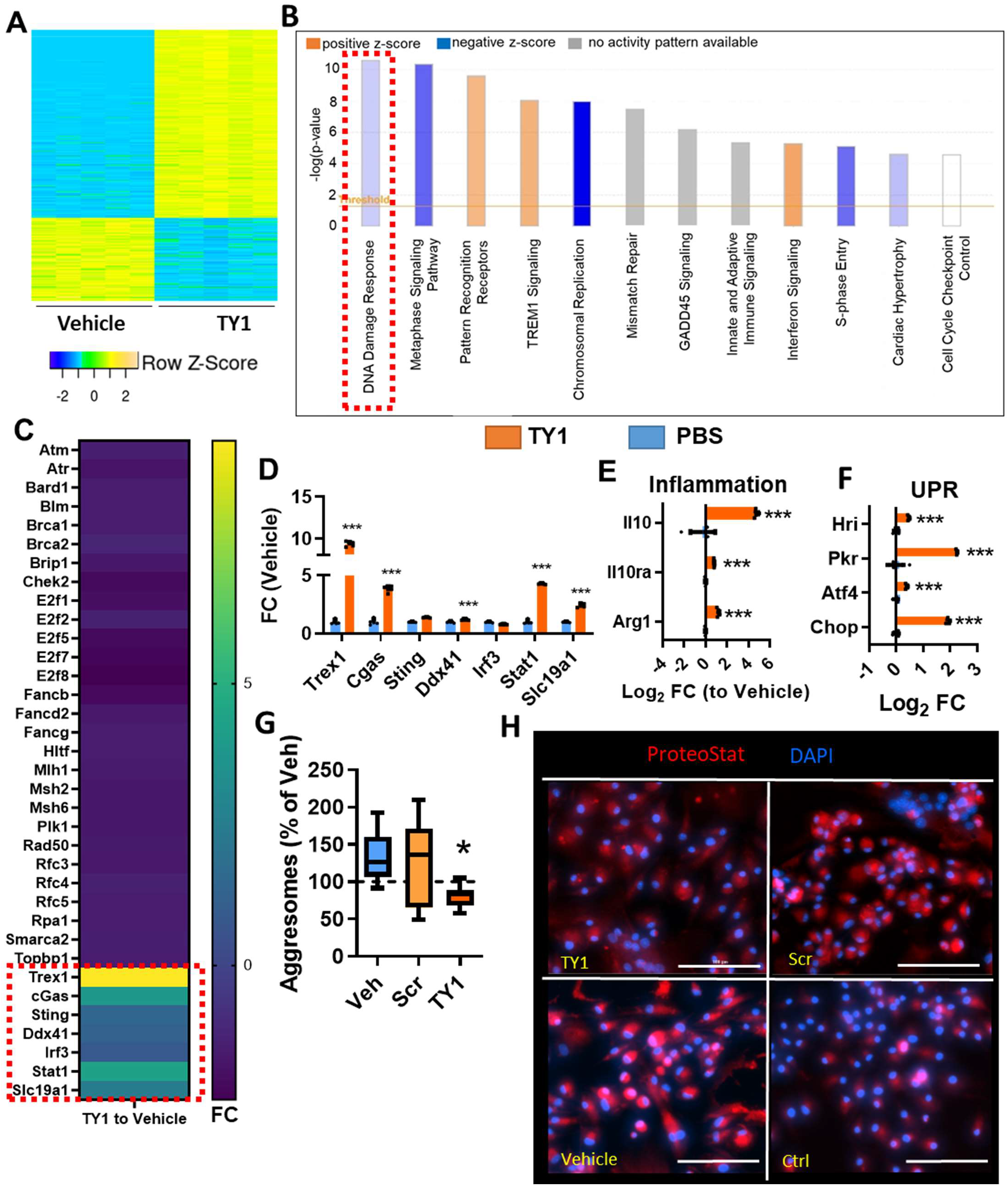
TY1 reduces cell stress through attenuation of genotoxicity. (**A**) Heatmap of RNA sequencing data from rat bone marrow-derived macrophages (BMDM) exposed to vehicle (PBS) or TY1 (n=4 biological replicates per group). (**B**) Ingenuity Pathway Analysis (IPA) identifies DNA repair as major implicated pathway in TY1-exposed macrophages. (**C, D**) Heatmap showing all differentially expressed DNA repair genes with unique enrichment of TREX1 signaling. Downregulation of inflammatory, (**E**) and unfolded protein response (UPR) genes (**F**). LPS-stressed macrophages exposed to TY1 showed a lower burden of misfolded protein aggregates (aggresomes) compared to scramble and vehicle as measured by the amyloid aggregate dye, ProteoStat (**G, H**; red; co-stained with DAPI, blue; n=7-8 biological replicates per group). Scale bars: 100 μm. Bars represent group means and error bars represent s.d. Analysis of two groups was done using a two-tailed Student’s t test with a 95% CI. Analysis of three or more groups was done using One-way ANOVA; *,P<0.05; **, P<0.01; ***P<0.001.

### TY1 attenuates cell stress through mRNA stabilization of TREX1

To investigate the mechanism whereby TY1 imparts its sweeping transcriptomic effects, we determined its subcellular localization at baseline and under stress conditions. In LPS-stressed BMDM, TY1 preferentially localizes to the nuclear fraction, in contrast to its predominantly cytosolic localization in unstressed BMDM (**fig. S2A**). To further examine proteins that associate with TY1, we performed a protein pull-down in which biotinylated TY1 (or the native NT4) was incubated with BMDM lysate followed by mass spectrometry of the bound fraction (**fig. S2B**). Both TY1 and NT4 associated with nuclear pore proteins (nucleoporins), shuttling proteins (Ran GTPase system), and mRNA binding proteins (**fig. S2C, D**).

Given that it localizes to the nucleus, we explored the possibility that TY1 might interact directly with selected transcripts. We thus performed mRNA pull down, using two complementary approaches: a “bait method” in which biotinylated TY1 is incubated with BMDM lysate, or a “probe method” exposing rat BMDM to TY1 followed by retrieval using a biotinylated anti-sense probe (**Fig. 3A**). Bound mRNA species retrieved by either approach were evaluated using RNA sequencing. The two methods yielded 99 mRNAs in common (**Fig. 3B**), which were then cross-referenced to previous sequencing data (**Fig. 2**). Of the 99 genes identified by both the bait and probe approaches, 14 were upregulated and 8 were downregulated (**Fig. 3C**). The gene with the greatest TY1-induced upregulation in rat BMDM was TREX1, i.e., 3’ repair exonuclease 1 (identified earlier in **Fig. 2C, D**), which was similarly upregulated in mouse macrophages (**fig. S3A**). In a mouse monocyte line (Raw264.7 cells), TY1 increased TREX1 protein expression (**fig. S3B, C**). Finally, Raw264.7 cells pre-incubated with TY1 were protected against DNA damage induced by serum starvation (**fig. S3D, E**). Both mRNA pulldown methods identified TREX1 as a potential binding partner of TY1 (**Fig. 3D, E**). Small RNAs can regulate mRNA transcripts through 3’ or 5’ untranslated regions (UTR)(*37*). A UTR-luciferase assay demonstrated preferential binding of TY1 to the 5’ UTR (**Fig. 3F**), but no binding to the 3’ UTR of human TREX1 (**Fig. 3G**), both compared to Scr. Furthermore, exposing BMDM to TY1 upregulated TREX1 expression (**Fig. 3H**). Binding of small RNAs to the 5’ UTR can enhance mRNA stability, loading to the ribosomal complex, and resistance to mRNA degradation(*38-40*). Alternatively, binding of the 5’ UTR could lead to enhanced transcription(*41*). Exposing Raw264.7 macrophages to the transcriptional inhibitor actinomycin D after exposure to TY1, Scr, or vehicle showed comparable mRNA lifetime curves for TREX1, excluding an mRNA stabilization effect of TY1 (**Fig. 3I**). TY1 partially rescued TREX1 expression (**Fig. 3J**) and Il10 expression (**Fig. 3K**) in BMDM transfected with TREX1-silencing RNA (siRNA). Therefore, TY1 upregulates TREX1 transcription by binding to the 5’ UTR and enhancing TREX1 expression. Direct overexpression of TREX1 (via knock-in plasmid; TREX1 OE) dramatically increased Il10 in BMDM (**Fig. 3L**), motivating us to test whether TREX1 upregulation might suffice to explain the salutary effects of TY1 in MI.

**Figure 3:**
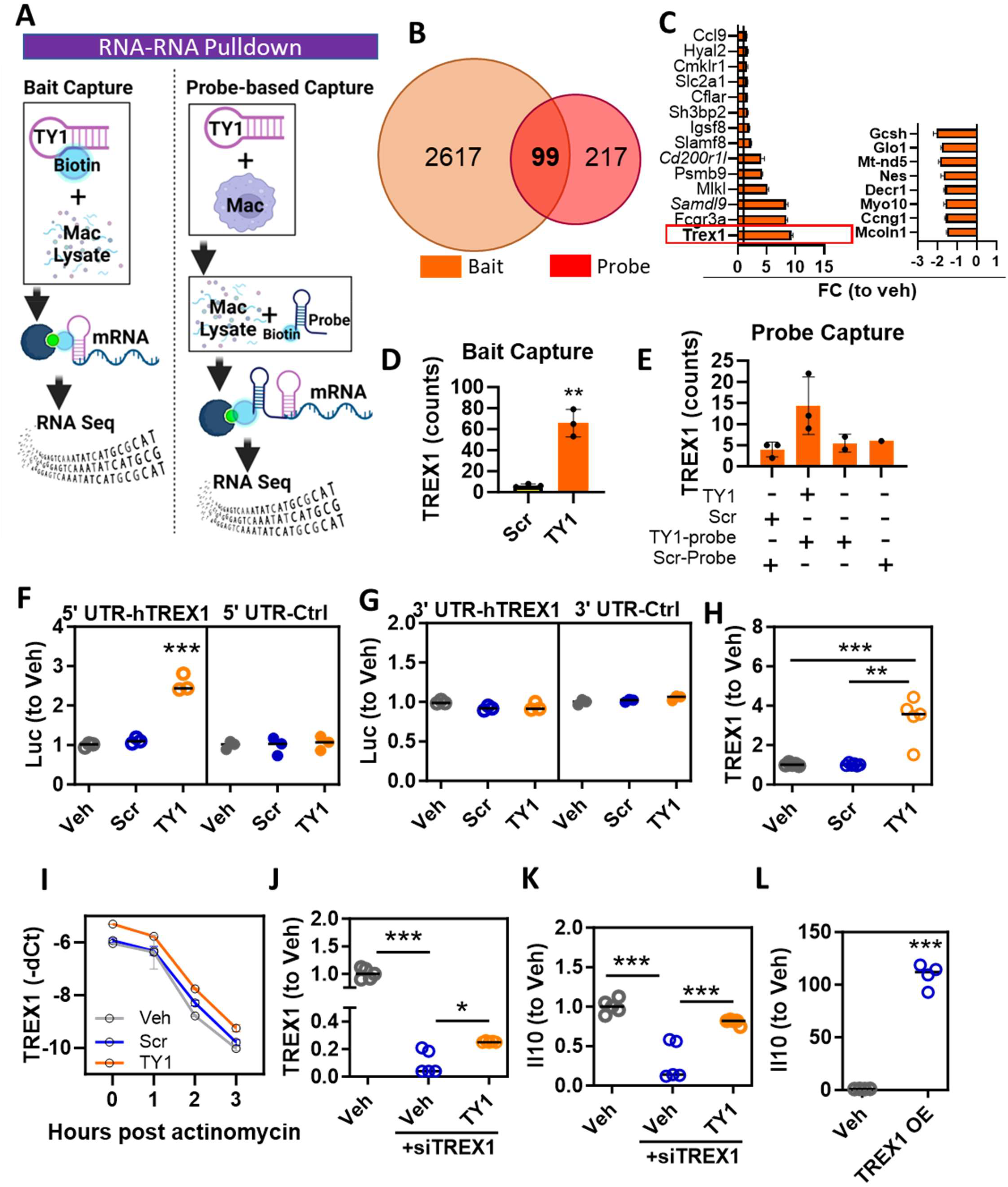
TY1 attenuates cell stress through mRNA stabilization of TREX1. (**A**) Schematic for two orthogonal approaches to identifying RNA targets of TY1. (**B**) Venn Diagram showing genes identified using TY1 RNA targets. (**C**) TY1 RNA targets up and down regulated in the RNA sequencing of macrophages exposed to TY1 (compared to macrophages exposed to vehicle; n=5 biological replicates per group). (**D, E**) Sequencing reads of RNA-RNA pulldown showing enrichment of TREX1 enrichment using the bait capture method (**D**) and the probe-capture method (**E**; n=3 biological replicated per group). Untranslated region (UTR)-luciferase assay demonstrating stabilization of luciferase signal via binding the 5’ UTR (**F**) but not the 3’ UTR (**G**) of TREX1. (**H**) QPCR validation of TREX1 upregulation in bone marrow-derived macrophages (n=3 biological replicates per group). (**I**) TY1 does not impact stability of existing TREX1 mRNA as shown by comparable decay rates (by qPCR) post treatment of Raw 264.7 macrophages incubated with the transcription inhibitor actinomycin D (10 μg/ml). Suppression of TREX1 in macrophages is partially rescued with TY1 exposure (**J**) along with Il10 expression, (**K**; n=4 biological replicates per group).(**L**) TREX1 upregulation leads to enhanced Il10 expression in macrophages (n=3 biological replicates per group). Bars represent group mean and error bars represent s.d. Significance was determined by one-way ANOVA with Tukey’s post-test; *P<0.05; **, P<0.01; ***, P<0.001.

### TY1 enhances tissue repair in MI

We began *in vivo* studies by investigating the toxicity of TY1. Healthy mice receiving twice-weekly intravenous (IV) injections of TY1 (0.15 mg/kg, based on prior dose optimization of EV-YF1(*23*), and again formulated in DharmaFECT®) for four weeks (**fig. S4A**) exhibited unchanged body weight (**fig. S4B**), blood counts (**fig. S4C**), organ weights (**fig. S4D**) and blood chemistry analyses (**fig. S4E**) relative to vehicle or Scr controls (also formulated with DharmaFECT®). Motivated by prior observations that EV-YF1(*21*) and NT4(*33*) are cardioprotective, we tested TY1 in two complementary MI models (**Fig. 4A**). In the first model, rats underwent coronary ligation for 45 minutes. Twenty minutes after reperfusion, they received IV TY1 (or Scr, each 0.15 mg/kg; or vehicle). Forty-eight hours post-MI, TY1 animals exhibited a decrease in infarct size (**Fig. 4B, C**) and lower circulating levels of cardiac troponin I, an ischemic biomarker (**Fig. 4D**), relative to vehicle or Scr. The decrease in infarct size directly demonstrates cardioprotection, while lower troponin I levels signify decreased cardiomyocyte necrosis. Consistent with TY1’s proposed mechanism of action, TREX1 protein levels were elevated (**Fig. 4E**), and DNA damage was attenuated (**Fig. 4F, G**), in the hearts of animals that had received TY1 (vs Scr or vehicle). Immunohistochemistry revealed no change in the overall levels of CD68 (a macrophage marker; **fig. S5A, B**), but increased TREX1 expression in macrophages (**fig. S5A, C**), in hearts of animals given TY1. Therapeutic benefits persisted at least 3 weeks post-MI (**Fig. 4H**), at which time rats that had received TY1 exhibited increased left ventricular function (**Fig. 4I, J**), attenuated chamber remodeling (**Fig. 4K, L**) and reduced myocardial scar (**Fig. 4M, N**) relative to vehicle. We then tested whether TY1’s cardioprotective effects could be reproduced in a clinically relevant porcine model of MI (**fig. S6A**). As in rats, IV infusion of TY1 post-MI in pigs reduced infarct size (**fig. S6B, C**). Taken together, the findings from two complementary MI models indicate that TY1 upregulates TREX1 (especially in infiltrating macrophages), reduces DNA damage, and augments cardiac tissue repair post-MI.

**Figure 4:**
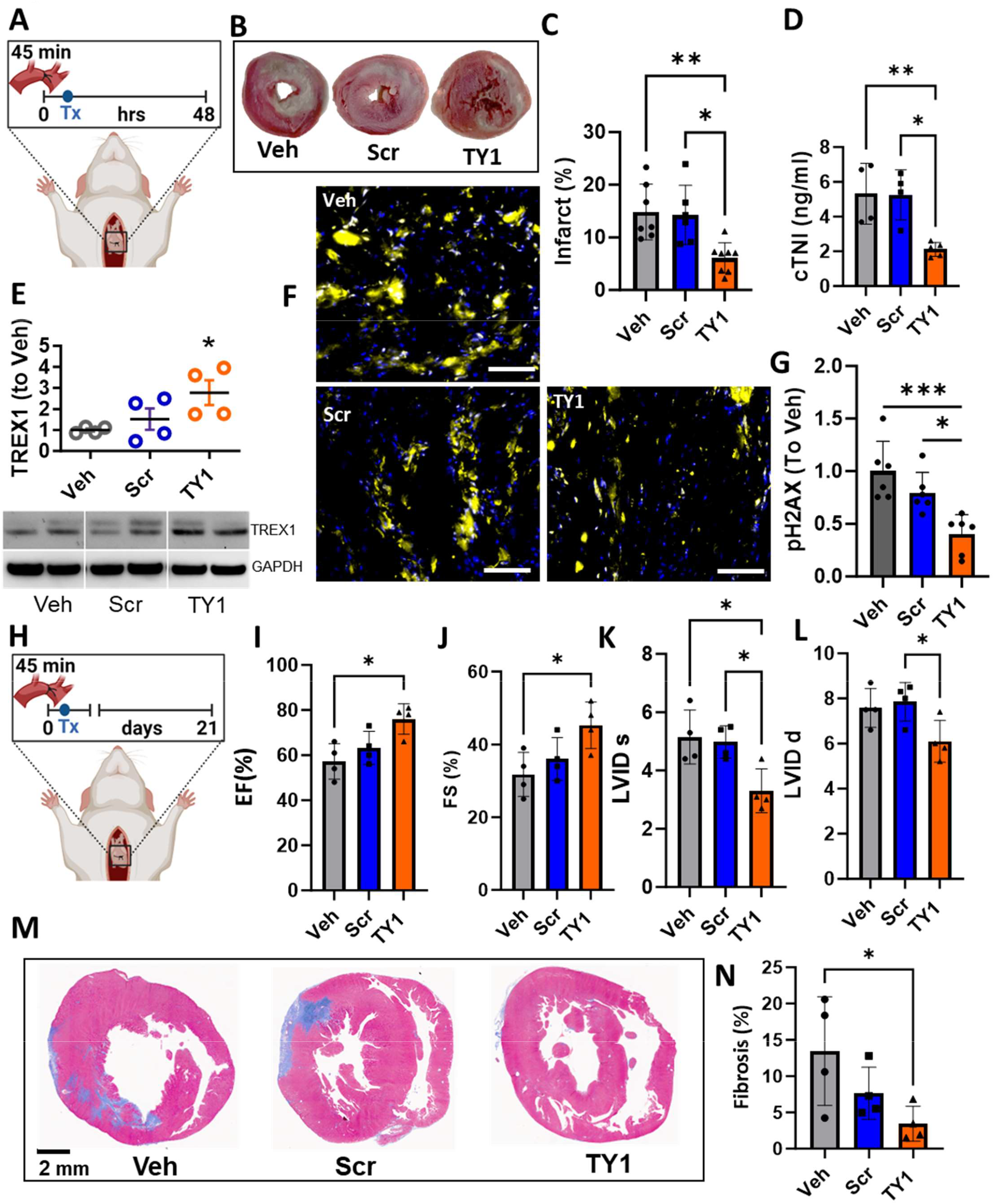
TY1 reduces infarct size in a rat model of MI. (**A**) Study design for TY1 administration in a rat model of acute myocardial infarction (n=4-5 animals per group). Rats received ischemia reperfusion injury (45 minutes of ischemia) followed by intravenous administration of TY1 (0.15 mg/kg), TY1 scramble control (0.15 mg/kg), or vehicle. Forty-eight hours post-injury, animals receiving TY1 had reduced infarct size (**B, C**) abd lower circulating cardiac troponin (**D**). (**E**) Western blot showing increase of TREX1 in LV tissue of animals receiving TY1 compared to scramble or control groups. (**F, G**) TY1-exposed animals had lower levels of genotoxic stress as measured by the DNA damage marker phosphor H2AX (pH2AX; red, scale bar: 1 mm) compared to vehicle and scramble groups. (**H**) Study design for long term follow-up of MI animals receiving a single dose of TY1, Scr, and vehicle. Rats receiving TY1 preserved ejection fraction (**I**), fractional shortening (**J**), end systolic (**K**) and diastolic volumes (**L**), and reduced fibrotic burden (**M, N**). Lines represent group mean and error bars represent s.d. Significance was determined by one-way ANOVA; *P<0.05; **P<0.01; ***P<0.001.

### Macrophages mediate the cardioprotective effects of TY1

The observation that TY1 upregulates TREX1 in cardiac macrophages led us to hypothesize that macrophages serve as effectors of TY1 benefits. Consistent with this hypothesis, TY1 cardioprotection was lost in macrophage-depleted animals (**Fig. 5A-C**). To test whether macrophages suffice to confer TY1 cardioprotection, we infused rat BMDM IV post-MI, and assessed infarct size. Before adoptive transfer, BMDM were manipulated by exposure to one of the following: TY1; vehicle; Scr; a TREX1 overexpression vector; or TY1 plus anti-TREX1 siRNA (siTREX1: **Fig. 5D**). Animals infused with macrophages exposed to TY1, or macrophages genetically enriched in TREX1, had smaller infarct sizes compared to Scr or vehicle (**Fig. 5E, F**).

**Figure 5:**
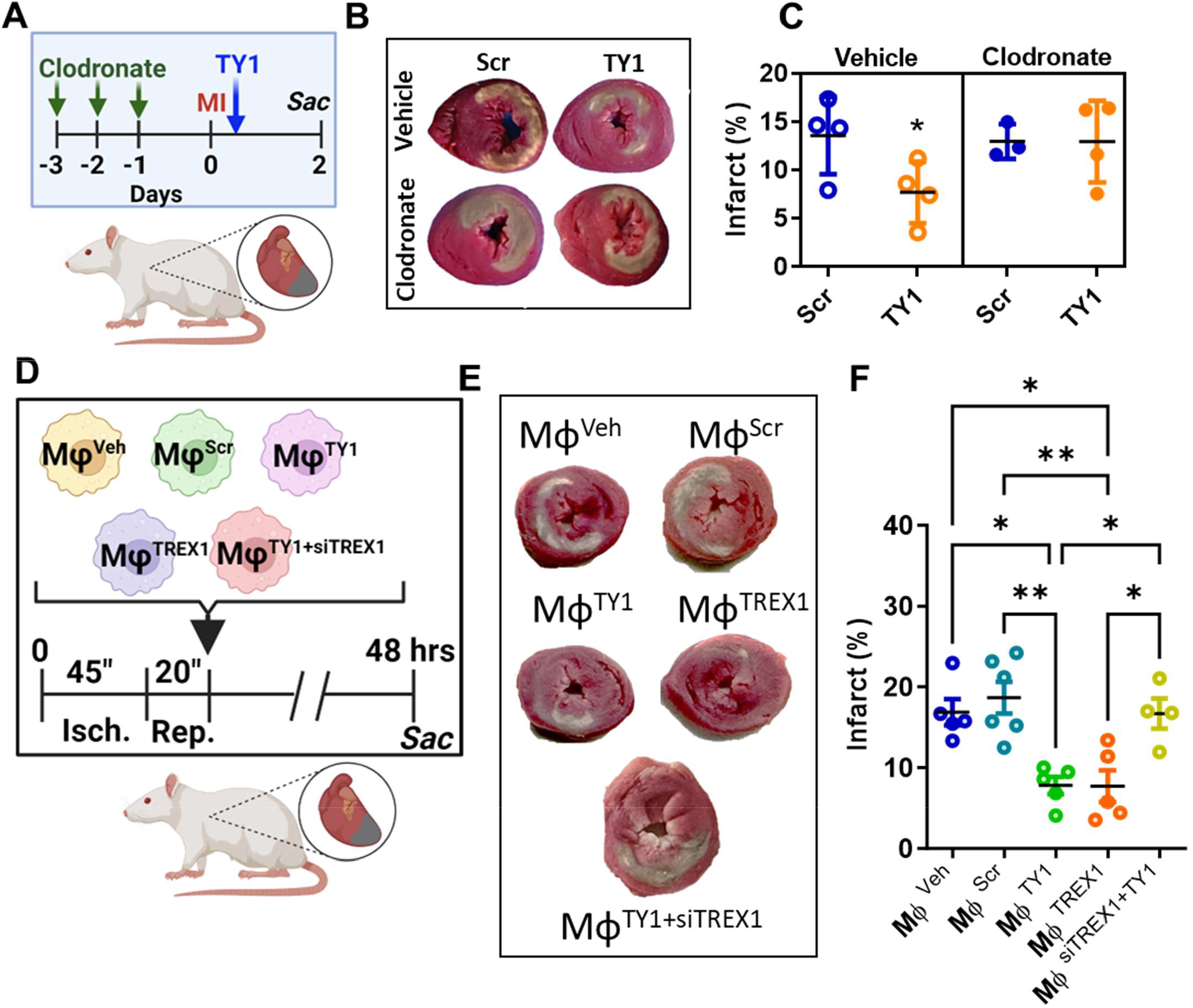
Macrophages mediate TY1 cardioprotection. (**A**) Macrophages were depleted from rats through daily intravenous administration of clodronate prior to induction of myocardial infarction. Animals were given TY1, scramble, or control 20 minutes post reperfusion. TY1 reduces scar size (compared to vehicle to scramble groups) in animals with functioning macrophages (**B, C**). However, TY1 therapeutic activity is attenuated by macrophage depletion in rats as shown by comparable scar size to vehicle or scramble groups (**B, C**; n=4 animals/group). In a separate study, rats with MI received tail vein infusions of macrophages (3× 10^6^ cells/animal) transfected with TY1, scramble, vehicle, or a TREX1 activating plasmid (**D**; n=3-5 animals per group). Animals with MI given macrophages overexpressing TY1 or TREX1 had significant reductions in scar size compared to animals given macrophages transfected with scramble or vehicle (**E, F**). Lines represent group mean and error bars represent s.d. Significance was determined by one-way ANOVA; *P<0.05; **P<0.01; ***P<0.001.

Thus, adoptive transfer of macrophages exposed to TY1 suffices to induce cardioprotection, an effect mimicked by adoptive transfer of macrophages overexpressing TREX1. The additional finding that siTREX1 blocked the cardioprotective effect of TY1 (**Fig. 5E, F**) directly links TREX1 upregulation in macrophages to the mechanism of TY1 action. We conclude that macrophages are necessary and sufficient for TY1 cardioprotection, which in turn is due to TREX1 upregulation.

## Discussion

DNA damage induced by oxidative stress, whether by acute injury (as in ischemia/reperfusion(*42*)), or during gradual and stochastic accumulation of genomic errors (as in aging(*43, 44*)), drives cell and tissue dysfunction. Nuclear DNA damage leads to DNA leakage into the cytosol, kickstarting cGAS-STING(*45*). TREX1 mitigates DNA damage, in part, by degrading cytosolic DNA and preventing cGAS-STING and downstream interferon gene cascades(*12, 46*). Here we demonstrate that a small synthetic ncRNA molecule, TY1, upregulates TREX1 and minimizes lethal cardiac injury post-MI. TY1 exerts its tissue-reparative effects by enhancing cells’ ability to relieve genotoxic stress. The benefits of TY1 are specific: a scrambled version had no effect, and the changes are directionally opposite to those expected from the nonspecific immunogenicity that plagues many other ncRNA drugs(*24*). Mechanistic dissection identified TREX1 as a target of TY1. By binding to the 5’ UTR of the TREX1 mRNA, TY1 enhances TREX1 translation and bioactivity. TREX1 degrades DNA fragments that would otherwise activate cGAS-STING and promote inflammation(*45*). Indeed, TREX1 deficiency leads to systemic hyperinflammation(*11*), manifested as systemic lupus erythematosus and other autoimmune disorders(*10, 47*).

We have found that TREX1 enhancement attenuates tissue injury associated with sterile inflammation in two models of MI, an effect mediated by macrophages. Depletion of macrophages abrogates the cardioprotective effects of TY1 *in vivo*; those effects are mimicked by adoptive transfer of macrophages overexpressing TREX1 and blocked by suppressing TREX1 in TY1-exposed macrophages. Thus, macrophages are both necessary and sufficient for TY1’s cardioprotective bioactivity *in vivo*. This finding is not entirely surprising, given that macrophages are known to mediate CDC-EV-induced cardioprotection(*16, 48, 49*), and TY1 was bioinspired by EV-YF1, an ncRNA that is especially plentiful in CDC-EVs(*21*). This sequence of insights—CDCs secrete EVs packed with obscure but bioactive ncRNA cargo—underlies the non-traditional development pathway for TY1: we first identified EV-YF1, demonstrated its disease-modifying bioactivity, then used the natural compound as a template for structural optimization, and finally probed mechanism. The discovery that TY1 upregulated TREX1 was not only serendipitous, but also fortuitous, enabling us to probe the DNA damage response with a trenchant new tool. The conventional drug development approach works in reverse, designing ncRNAs to target a particular predetermined disease pathway. Reflecting the unprecedented development paradigm, TY1 does not work as a microRNA or small interfering RNA, nor is it an aptamer or antisense molecule. Instead, TY1 is the prototype—both structurally and conceptually—for a new class of therapeutic candidates.

Our results validate TREX1 upregulation as a therapeutic principle in MI, a common inflammatory disorder of immense public health significance(*50*). Despite nearly a century of investigation(*51*), the only strategy proven to reduce infarct size in patients remains prompt reperfusion(*52*). Here we have identified DNA damage as a novel target for cardioprotective strategies. Though we focused on MI, the same cell stress pathways figure prominently in diverse pathologies. Indeed, we have reported that TY1 has disease-modifying bioactivity in models of heart failure with preserved ejection fraction(*53*) and scleroderma(*54*), two conditions refractory to treatment in which inflammation and fibrosis feature prominently. TY1 is predictably worth testing in models of other diseases driven by DNA damage and/or inflammation, including lupus(*55, 56*) and Duchenne muscular dystrophy(*57, 58*). More generally, TY1 will allow us to probe the role of DNA damage responses even in settings where their involvement has been previously unsuspected. Such speculation seems plausible given that TY1’s targets are biologically fundamental and highly conserved: TREX1 has bacterial homologs(*59*), while the cGAS/STING immune recognition pathway dates back >600 million years to the Precambrian era(*60*). This enduring conservation attests to the critical importance of genotoxic stress in shaping physiological and pathological cellular responses.

## Acknowledgments

We thank Liang Li for help with animal studies, Jeanna Huynh for assistance with the manuscript, and the Cedars-Sinai Genomics Core for RNA sequencing services. We also thank Dr. Yu Chen of the UCLA Molecular Instrumentation facility for mass spectrometry services. Figure schematics were generated using Biorender.com

## Funding

National Heart Lung and Blood Institute grant R01 HL164588 and T32 HL116273 (EM) National Heart Lung and Blood Institute grant R01 HL142579 (AGI)

### Author contributions

Conceptualization: EM and AGI

Methodology: AGI, KM, AC, XJ, RR, JD, and AR

Investigation: KM, AC, XJ, RR, JD, AR, CL, KL, SY, AM, KT, AN, JL, CHM, YH, NM, and ANK

Funding Acquisition: EM and AGI

Project Administration: EM and AGI

Supervision: EM, AGI, RR, and JD

Writing Original Draft: AGI

Writing-review and editing: EM, AGI

### Data and materials availability

All data are available in the main text or the supplementary materials.

## Supplementary Materials

Materials and Methods

Supplementary Text Figs.

S1 to S6

Tables S1 to S1

**Graphical abstract.**
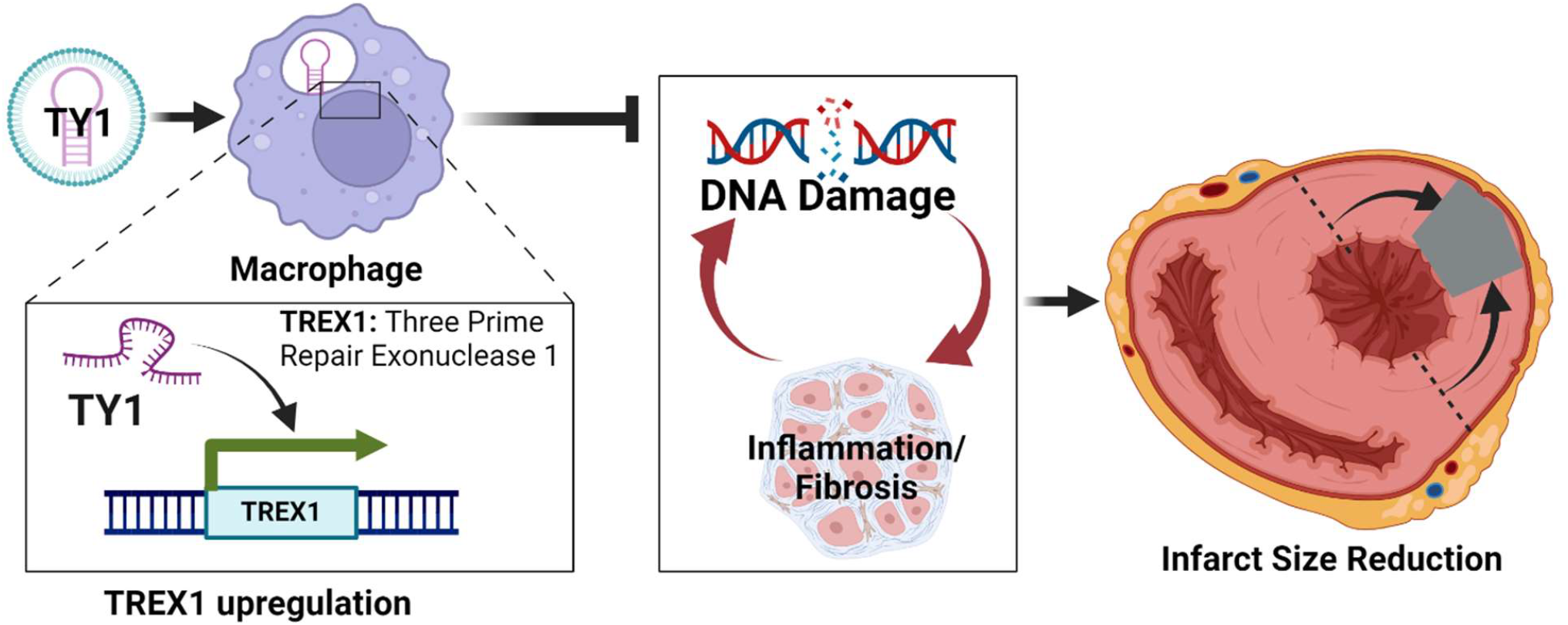
Based on naturally-occurring small Y RNAs, we developed TY1, a new chemical entity with sweeping impact on innate immunity and DNA damage pathways in macrophages. Mechanistic dissection reveals TY1 binds to and enhances expression of TREX1, a key regulator of genotoxic stress. *In vivo*, TY1 attenuates cardiac damage induced by ischemia/reperfusion in myocardial infarction. Upregulation of TREX1 in macrophages is sufficient and necessary for TY1’s disease-modifying bioactivity.

