## Supplementary Material for "Augmentation of DNA exonuclease TREX1 in macrophages as a therapy for cardiac ischemic injury"

**The PDF file includes:**

Materials and Methods
Supplementary Text
Figs. S1 to S6
Tables S1 to S1

### Materials and Methods

*Macrophage Drug Screen*

***Synthesis and formulation of RNA compounds*** TY1 and derivatives were synthesized (IDT) and mixed with 3 µl DharmaFECT® transfection reagent (Horizon Discovery) to a final volume of 100 µl in serum-free media. Following agitation and incubation (to form liposomal-RNA complexes), the preparation is added to macrophage culture media to a final concentration of 80 nM, for *in vitro* applications. For *in vivo* applications (intravenous), a similar formulation is prepared to achieve a dose of 0.15 mg/kg. The same process is repeated with each of the new chemical variants of TY1 and with Scramble (Scr).

***Rodent bone marrow cell isolation, macrophage differentiation, and activation*** Femurs were isolated from 7-10-week-old Wistar Kyoto rats or C57BL6 mice, as indicated. Bone marrow was isolated by flushing with phosphate-buffered saline (PBS) and then filtering through a 70 µm mesh. Red blood cells were lysed with ACK buffer (A1049201, Invitrogen); the remaining cells were resuspended in IMDM (Gibco) containing 20 ng/ml M-CSF (RP8643, Fisher Scientific) and plated. The media was exchanged every 2-3 days until day 7, in a process that yields bone marrow-derived macrophages (BMDM). BMDM were transfected with small Y RNAs (or Scr; 80 nM) after encapsulation in DharmaFECT®. For lipopolysaccharide (LPS) exposure, macrophages were pretreated with vehicle (DharmaFECT® + PBS), TY1 (80 nM, in DharmaFECT® + PBS), or TY1 Scr control (80 nM, in DharmaFECT® + PBS) for three hours followed by exposure to LPS (10 ng/ml; Cayman Chemical). Twenty-four hours after exposure to each test agent, RNA was extracted and subjected to RNA seq (detailed below) or to analysis by quantitative polymerase chain reaction (qPCR) to determine transcript levels, as indicated.

***Human bone marrow-derived macrophages*** Human peripheral blood monocytes were purchased (STEMCELL Technologies), and cultured in RPMI 1640 Medium with 10% FBS, and human M-CSF (Invitrogen) to create macrophages. Macrophages were transfected with small Y RNAs (or Scr; 80 nM) after encapsulation in DharmaFECT®. Twenty-four hours after exposure to each test agent, RNA was extracted and subjected to analysis by quantitative polymerase chain reaction (qPCR) to determine transcript levels of the anti-inflammatory cytokine IL10, and HPRT1, a housekeeping enzyme used as an endogenous control.

***Comet Assay***

BMDM cultured in IMDM and 2% FBS were transfected with TY1, vehicle (transfection agent [Dharmafect] only) or control (untreated) for 4-5 hours in Optimem. Following transfection, the media was changed to fresh IMDM with 2% FBS overnight. Raw264.7 cells cultured in DMEM (ATCC) and 2% FBS were transfected with TY1, scramble, or control (untreated) for 4-5 hours in Optimem. Following transfection, the media was changed to fresh DMEM with 2% FBS overnight. Cells were washed twice with PBS, then incubated with CTS Versene (Gibco) for 5 minutes at 37°C and gently scraped with a cell scraper to detach cells. Cells were centrifuged at 700 x g for 2 minutes, supernatant discarded, and the cell pellet was washed once with cold PBS. Cells were centrifuged again and resuspended in 25µL of cold PBS. Cells were frozen at -80°C to permeabilize cell membrane. Positive control was made using healthy cells that were serum starved on ice for 6 hours. Following permeabilization, cells were resuspended to a concentration between 1 x 105 cells/mL to 5 x 105 cells/mL. Cells were combined with low-melting point Comet agarose in a ratio of 1:10 by volume and pipetted onto pre-prepared microscope slides with 1% agarose. Glass coverslips were used to set the agarose in place. Slides were transferred to the 4°C for 15 minutes in the dark before removing coverslips. Slides were allowed to fully set at 4°C for an additional 15 minutes. Slides were immersed in cold Lysis Buffer (2.5M NaCl, 100mM EDTA, 10% DMSO, 10% 10X Lysis Solution from Cell Biolabs, pH 10.0) for 60 minutes at 4°C in the dark to further permeabilize cells. Lysis Buffer was removed, and slides were immersed in cold Alkaline Solution (300mM NaOH, 1mM EDTA) for 30 minutes at 4°C to denature DNA. Alkaline Solution was removed, and slides were transferred into a horizontal cold electrophoresis chamber with cold Alkaline Electrophoresis Solution (300mM NaOH, 1mM EDTA). Voltage was applied for 45 minutes at 1 volt/cm with a current setting of 300 mA. Slides were removed from the electrophoresis chamber and washed 3X with pre-chilled DI H2O for 2 minutes per wash. Slides were washed with cold 70% ethanol for 5 minutes. Slides were allowed to dry at 37°C for 30 minutes. Slides were stained with diluted Vista Green DNA Dye from Cell Biolabs (in TE Buffer, pH 7.5) for 15 minutes at room temperature, then left to dry before viewing with Cytation 5 using the GFP FITC filter. Comets were quantified using the "Cellular Analysis" program in Cytation 5 to measure the size, area, and fluorescence of the comet head and tail to generate the Olive Tail Moment calculation.

***RNA isolation and quantitative PCR*** Total RNA was extracted from mouse, rat, and human cells using RNeasy Plus Kit (74136, QIAGEN) and MaXtract High Density (129056, QIAGEN) and/or miRNeasy Mini Kit (217004, QIAGEN). cDNA was synthesized from RNA using High-capacity cDNA Reverse Transcription Kit (4368813, Applied Biosystems) according to the manufacturer's protocol. Real-time PCR (QuantStudio 12K Flex Real-Time PCR system; Thermo Fisher Scientific) was performed in triplicate using the following TaqMan Gene Expression Assay probes (Cat# 4331182; with corresponding Assay IDs in Supplementary Table 1); Differential gene expression analysis was done using the ΔΔCt method.

***RNA Sequencing*** Cell and tissue RNA samples were sequenced at the Cedars-Sinai Genomics Core as described(*61*). Total RNA samples were assessed for concentration using a Qubit fluorometer (ThermoFisher Scientific, Waltham, MA) and for quality using the 2100 Bioanalyzer (Agilent Technologies, Santa Clara, CA). Library construction was performed using the QIASeq Stranded RNA Library kit (Qiagen, Hilden, Germany) with QIAseq FastSelect - rRNA HMR Kit (Qiagen) for ribosomal RNA depletion. Library concentration was measured with a Qubit fluorometer and library size on a Bioanalyzer. Libraries were multiplexed and sequenced on a NovaSeq 6000 (Illumina, San Diego, CA) using 75 bp single-end sequencing. On average, approximately 50 million reads were generated from each sample.

***Data analysis*** Raw sequencing data were demultiplexed and converted to FASTQ format by using bcl2fastq v2.20 (Illumina, San Diego, California). Reads were aligned to the GRCm38 reference genome (http://www.gencodegenes.org) using STAR (version 2.6.1)4 with default parameters. Gene expression was quantified by RSEM (version 1.2.28)5 to generate a raw count expression matrix with gene identities as rows and samples as columns. DESeq2 (version 1.26.0)6 was used to normalize the raw count expression and correct the batch effect.

***Intracellular protein aggresome assay*** BMDM were pre-exposed to TY1 or Scr for 4 hours, then exposed to LPS (10 ng/ml) overnight. After washing twice with PBS, cells were fixed in PFA 4% for 15 minutes, permeabilized, and incubated with PROTEOSTAT dye (1:10,000 dilution) for 30 min at RT (ENZ-51035-K100, Enzo Life Sciences). PROTEOSTAT-stained cells were imaged by epi-fluorescence microscopy for aggresomes and aggresome-like inclusion bodies.

***Nuclear/cytosolic RNA isolation and qPCR analysis*** Nuclear and cytosolic RNA were purified using Cytoplasmic and Nuclear RNA Purification Kit (Norgen Biotek) according to the manufacturer's protocol. Reverse transcription was performed using TaqMan® microRNA Reverse Transcription Kit (Applied Biosystems) per the manufacturer's protocol, using specific primers for TY1. Real-time PCR was performed using TaqMan Fast Advanced Master Mix and appropriate TaqMan Gene Expression Assay (Thermo Fisher Scientific). The reaction was performed in QuantStudioTM 12K Flex Real-Time PCR System, and each reaction was performed in triplicate samples and adjusted using snu6 as housekeeping (Life Technologies). Cycling conditions were performed according to the TaqMan protocol. Where appropriate, the $2^{-\Delta\Delta Ct}$ method was used to determine gene expression fold change.

***Protein Pulldown*** TY1 and Scr were labeled with desthiobiotinylated cytidine bisphosphate at the 3´ end of the RNA strands using T4 RNA ligase (Thermo Scientific Pierce RNA 3´ End Desthiobiotinylation Kit). Labeled RNA was captured using 50 µL streptavidin magnetic beads in

RNA Capture Buffer for 30 minutes at room temperature. Beads were washed twice in 20mM Tris (pH 7.5), once in Protein-RNA Binding Buffer, and 400 µg of BMDM extract was added per sample. Samples were incubated for 2 hours at 4°C, washed three times with Wash Buffer, and eluted after 15 minutes of incubation at 37°C with Biotin Elution Buffer. Samples were then analyzed by mass spectrometry (Molecular Instrumentation Center, UCLA).

*RNA-RNA Pulldown*

***Bait capture method***: To identify mRNA targets of TY1 using bait capture, lysates (400 µg of protein extract per sample) were incubated with 3' biotinylated TY1 and Scr sequences. After samples were washed with binding and wash buffers to remove unbound material (using a magnetic stand), RNA was extracted using column-based RNA purification (miRNAeasy; Qiagen). Following RNA purification, concentrations were measured, standardized, and submitted to the Cedars-Sinai Genomics Core for RNA quality evaluation (to ensure RNA integrity number [RIN]) followed by total RNA sequencing.

***Probe-based mRNA capture***: BMDM were incubated with 80 nM TY1, Scr, or vehicle for 18 hours. Cell lysates were then obtained and incubated for 18 hours with probe sequences relevant to the group (i.e., TY1 probes for TY1-exposed samples; Scr-specific probes for Scr-exposed samples; and both probe types for Vehicle groups, as negative controls; 100 pmol per sample). RNA was then extracted and sequenced as described in the bait-capture method above.

***3' and 5' UTR binding assays***

***3' UTR binding assay:*** HT1080 cells were cultured in IMDM with 10% FBS and seeded at 60-80% confluence prior to transfection. Cells were transfected with plasmids at a concentration of 120 ng/well (in FuGene HD transfection reagent [Promega] at a ratio 5:1 Fugene:vector) containing constructs containing the 3' UTR sequence of human TREX1 (or respective controls; Gene Copoeia) upstream of a Gaussia luciferase reporter constructs for 4 hours. Cells were then transfected with 80 nM of TY1, Scr, or vehicle formulated in DharmaFECT® for 15 hours. Conditioned media from wells was then collected and measured for luminescence emission using Secrete-Pair Dual Luminescence Assay Kit (gene Copoeia) using a microplate reader.

***5' UTR binding assay:*** HEK293T cells were cultured in EMEM with 10% FBS and seeded at 60-80% confluence prior to transfection. Cells were transfected with plasmids at a concentration of 120 ng/well (in DNAfectin 2100 transfection reagent [ABM]) containing constructs containing the 5' UTR sequence of human TREX1 (or respective controls; ABM, Inc.) upstream of a firefly luciferase reporter constructs for 4 hours. Media was then replaced to complete media overnight and the cells were then transfected with 80 nM of TY1, Scr, or vehicle formulated in DharmaFECT® for 5 hours. Wells were then measured for luminescence emission using a microplate reader the day after using Luciferase assay kit (ABM).

***Transcription inhibition assay***

Raw 264.7 cells (ATCC) were cultured at 80% confluence in DMEM in 2% FBS and transfected with vehicle (DharmaFECT® only) or 80 nM of scramble or TY1 for 4 hours. Media was then replaced with fresh media containing 10µg/ml of Actinomycin D (in DMSO; Sigma Aldrich). RNA was collected from cells at time 0, 1 hour, 2 hours, and three hours post-exposure with Actinomycin to measure the decay of existing TREX1 mRNA.

***Experimental Animals*** All studies were performed at Cedars-Sinai Medical Center in accordance with the Institutional Animal Care and Use Committee guidelines.

***Rat Ischemia/Reperfusion Model*** Rats (7-10-week-old female Wistar Kyoto; Charles River Labs, Wilmington, MA) were housed in a pathogen-free facility (cage bedding: Sani-Chips, PJ Murphy) with a 14 hours/10 hours light/dark cycle with food (PicoLab Rodent Diet 20 [no. 5053], Lab Diet) and water provided ad libitum. To induce MI, anesthetized rats underwent thoracotomy at the fourth intercostal space to expose the heart. A 7–0 silk suture was used to ligate the left anterior descending (LAD) coronary artery for 45 min, then removed to allow reperfusion. Twenty min later, vehicle (PBS with DharmaFECT®), TY1, or Scr (at 0.15 mg/kg, formulated in DharmaFECT® as described above) was injected into the retro-orbital sinus.

***Macrophage depletion using clodronate liposomes*** To deplete macrophages, rats were pre-infused with 1 mL clodronate liposomes (Liposoma) via tail vein infusion daily for 3 days prior to induction of MI(*48*).

***Adoptive transfer of macrophages*** Rat BMDM (prepared as described above) were incubated in vehicle (DharmaFECT® + PBS), 80 nM TY1 or 80 nM Scr (formulated in DharmaFECT®), or transfected a TREX1 activation plasmid formulated in FuGENE HD (Promega) transfection reagent for 24 hours. In the MI model, 20 minutes after reperfusion, each animal received a single tail vein infusion of $3x10^6$ BMDM which had been exposed to TY1, Scr, or vehicle for 24 hours, as described.

***Rat infarct size measurement*** Two days post-MI, 10% KCl was injected into the LV cavity; hearts were harvested, washed in PBS, and cut into 1-mm sections from apex to base, above the infarct zone. Sections were incubated with a 1% solution of 2,3,5-triphenyl-2H-tetrazolium chloride (TTC, Sigma-Aldrich) for 30 minutes at 37°C in the dark and washed with PBS. Then, sections were imaged and weighed. The infarcted zones (white) were delineated from viable tissue (red) and analyzed (ImageJ software). Infarct mass was calculated in the tissue sections according to the following formula: (infarct area/tot area)/weight (g).

***Cardiac troponin I ELISA*** Blood was collected from animals at 24 hours (from the tail vein) or study endpoint (from the LV cavity) in EDTA tubes. After incubation at 4°C for 30 minutes, plasma was obtained by 15-minute centrifugation at 4000 rpm. Cardiac troponin I was quantified using the rat cardiac troponin-I ELISA kit (Life Diagnostics) according to the manufacturer's protocol.

***Immunohistochemistry*** Tissues were embedded in OCT compound and frozen in 2-methyl butane precooled in liquid nitrogen, then stored at –80°C until sectioning. Serial sections of the heart were cut at the mid-papillary level in the transverse plane to 6 µm using a cryostat (CM3050S, Leica) and adhered to superfrost microscope slides. Cryosections of the heart were fixed with 4% paraformaldehyde solution (Fisher Scientific, AAJ19943K2) for 10 minutes, washed with cold PBS, permeabilized with 0.25% Triton™ X-100 (Millipore Sigma, T8787), and blocked (Protein Block, Dako with 0.05% Saponin [Sigma-Aldrich, S4521]) for 30 minutes at room temperature. Following the 30-minute block, slides were incubated overnight with primary antibodies diluted in a blocking solution at 4°C. Primary antibodies are as follows: CD68 (Abcam ab125212), TREX1 (NBP1-76977, Novus Biologicals), and pH2AX (D27C4, Cell Signaling Technologies). After the overnight incubation, slides were washed with PBS (3 times 5 minutes) and incubated with the appropriate Alexa Fluor–conjugated secondary antibody (1:500, Invitrogen) for 2 hours at room temperature. Following the secondary incubation, slides were washed with PBS (3 times 10 minutes) and coverslips were mounted with Fluoroshield with DAPI (Sigma-Aldrich) mounting medium. Slides were imaged using fluorescence microscopy (Cytation 5, Biotek) and quantified using Cytation 5 or ImageJ software. Pooled data were generated by averaging data from at least 4 fields per heart section.

***Porcine Ischemia/Reperfusion Model*** Adult female Yucatan mini-pigs were premedicated with ketamine 20 mg/kg IM, atropine 0.05 mg/kg IM, and acepromazine 0.25 mg/ kg IM. Animals were subsequently induced with propofol 2-4 mg/kg IV to effect, intubated, and maintained on isoflurane 2-3%. Amiodarone 10 mg/kg IV loading dose, then 0.5 mg/kg IV as needed, and lidocaine 2 mg/kg/min were given for ventricular arrhythmias. Heparin 100 IU/kg IV was given for anticoagulation. By inflating the balloon of an angioplasty catheter placed fluoroscopically, the proximal left anterior descending artery was occluded for 90 minutes, after which the balloon was deflated to allow reperfusion. Twenty ($\pm$10) minutes post-MI, animals received an IV dose of 0.15 mg/kg TY1 or Scr (formulated in DharmaFECT® + PBS) or vehicle (DharmaFECT® + PBS) in a total volume of 5 ml. Blood samples were acquired before balloon occlusion, after reperfusion, and at the study endpoint. Two days post-MI, pigs underwent induction of anesthesia and a lateral thoracotomy. An angioplasty balloon was placed at the level of the original occlusion during the MI procedure. With the balloon deflated, animals were infused with Thioflavin T (50 ml, 2% solution diluted with PBS) by direct injection into the left atrium over 60 seconds. The angioplasty balloon was then inflated and animals were infused with gentian violet (50 ml, 1.6% solution diluted with 40 ml PBS and 10 ml ethanol) by direct injection into the left atrium over 60 seconds. Finally, mini-pigs were sacrificed, and the heart was explanted and sectioned into 1 cm thick short-axis slices. Gentian violet stains the non-ischemic zone blue; the region of the heart that is not perfused by gentian violet represents the ischemic area at risk (AAR). Thioflavin T is a fluorescent dye (visualized under UV light), which stains endothelium receiving blood flow. Then, transverse ventricular slices were incubated with 2% TTC for 20 min at 37°C. TTC stains viable myocardium brick red, while the infarct appears white/yellow.

***Statistical analysis*** Statistical parameters including the number of samples (n), descriptive statistics (mean and standard deviation), and significance are reported in the figures and figure legends. Differences between groups were examined for statistical significance using the Student's t-tests or analysis of variance with Tukey's post hoc test. Differences with $p$ values < 0.05 were regarded as significant.

### Supplementary Text

Subhead

Type or paste text here. This should be additional explanatory text, such as: extended technical descriptions of results, full details of mathematical models, extended lists of acknowledgments, etc. It should not be additional discussion, analysis, interpretation, or critique.

Fig. S1.

### Supplemental Figure 1: TY1 attenuates cell stress signaling in macrophages

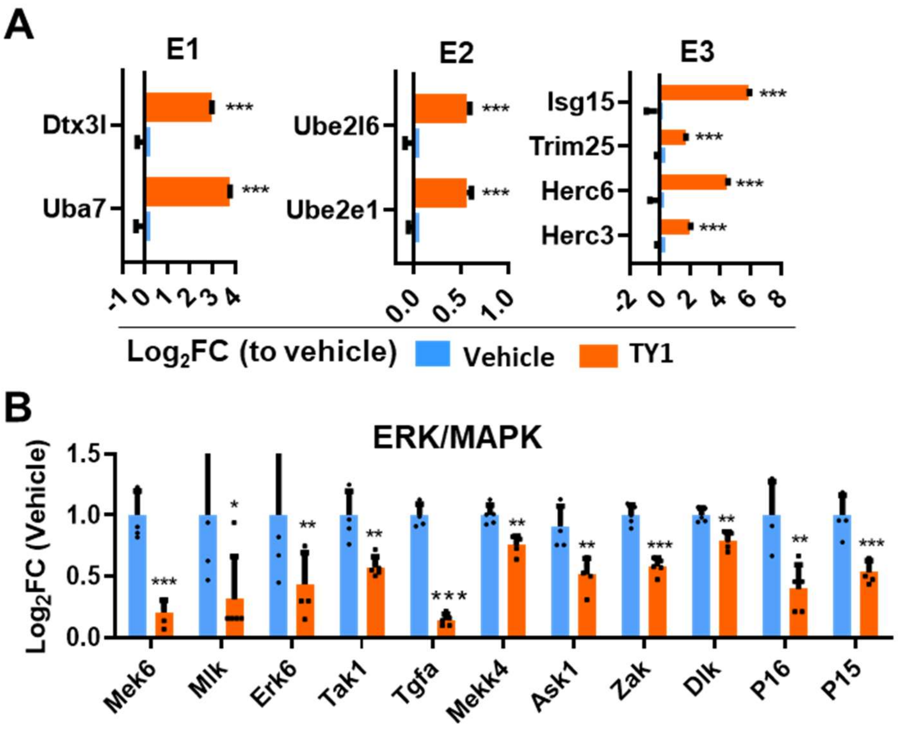


**Supplemental Figure 1: TY1 attenuates cell stress signaling in macrophages.**
(**A**) In bone marrow-derived macrophages upregulated genes of the ubiquitination pathway and suppressed stress-induced MAP-ERK signaling (**B**).

**Fig. S2.**

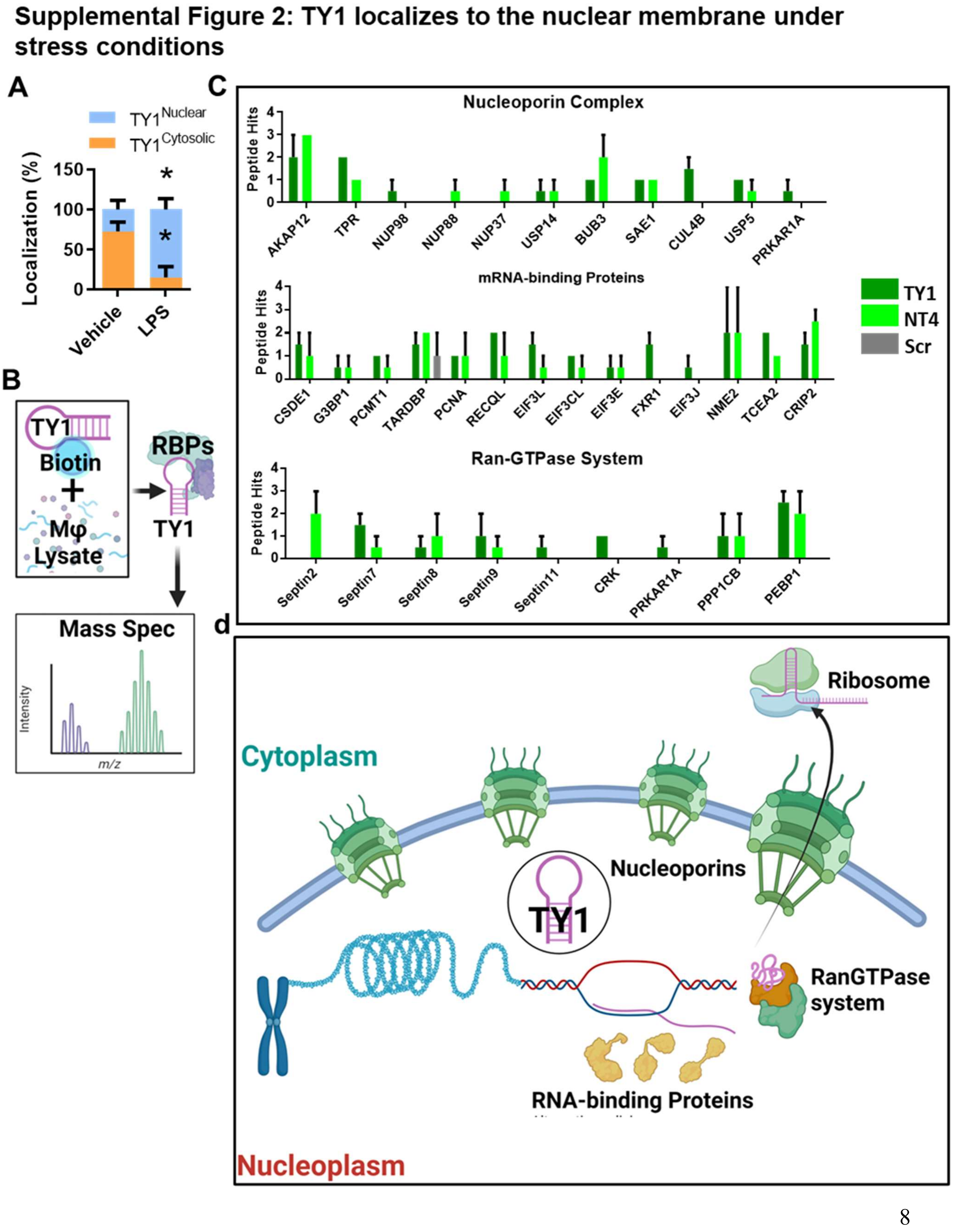

Supplemental Figure 2: TY1 localizes to the nuclear membrane under stress conditions
A
TY1 Nuclear
TY1 Cytosolic
Localization (%)
150
100
50
0
*
*
Vehicle
LPS
B
TY1
Biotin
+
Mφ
Lysate
RBPs
TY1
Mass Spec
Intensity
m/z
C
Nucleoporin Complex
Peptide Hits
AKAP12
TPR
NUP98
NUP88
NUP37
USP14
BUB3
SAE1
CUL4B
USP5
PRKAR1A
mRNA-binding Proteins
CSDE1
G3BP1
PCMT1
TARDBP
PCNA
RECQL
EIF3L
EIF3CL
EIF3E
FXR1
EIF3J
NME2
TCEA2
CRIP2
TY1
NT4
Scr
Ran-GTPase System
Septin2
Septin7
Septin8
Septin9
Septin11
CRK
PRKAR1A
PPP1CB
PEBP1
d
Ribosome
Cytoplasm
Nucleoporins
TY1
RanGTPase system
RNA-binding Proteins
Nucleoplasm

**Supplemental Figure 2 TY1 localizes to the nuclear membrane under stress conditions.**

(**A**) Enrichment of TY1 in the nuclear fraction of LPS-stimulated macrophages compared to vehicle-stimulated macrophages (n= 3 biological replicates per group). (**B**) Mass Spectrometry of protein pull down from incubating macrophage lysates with biotinylated TY1 demonstrates association with nucleoporin proteins, RAN GTPase proteins, and mRNA binding proteins (**C**). Collectively, these finding suggest that TY1 associates with proteins of the inner nuclear membrane (**D**).

Fig. S3.

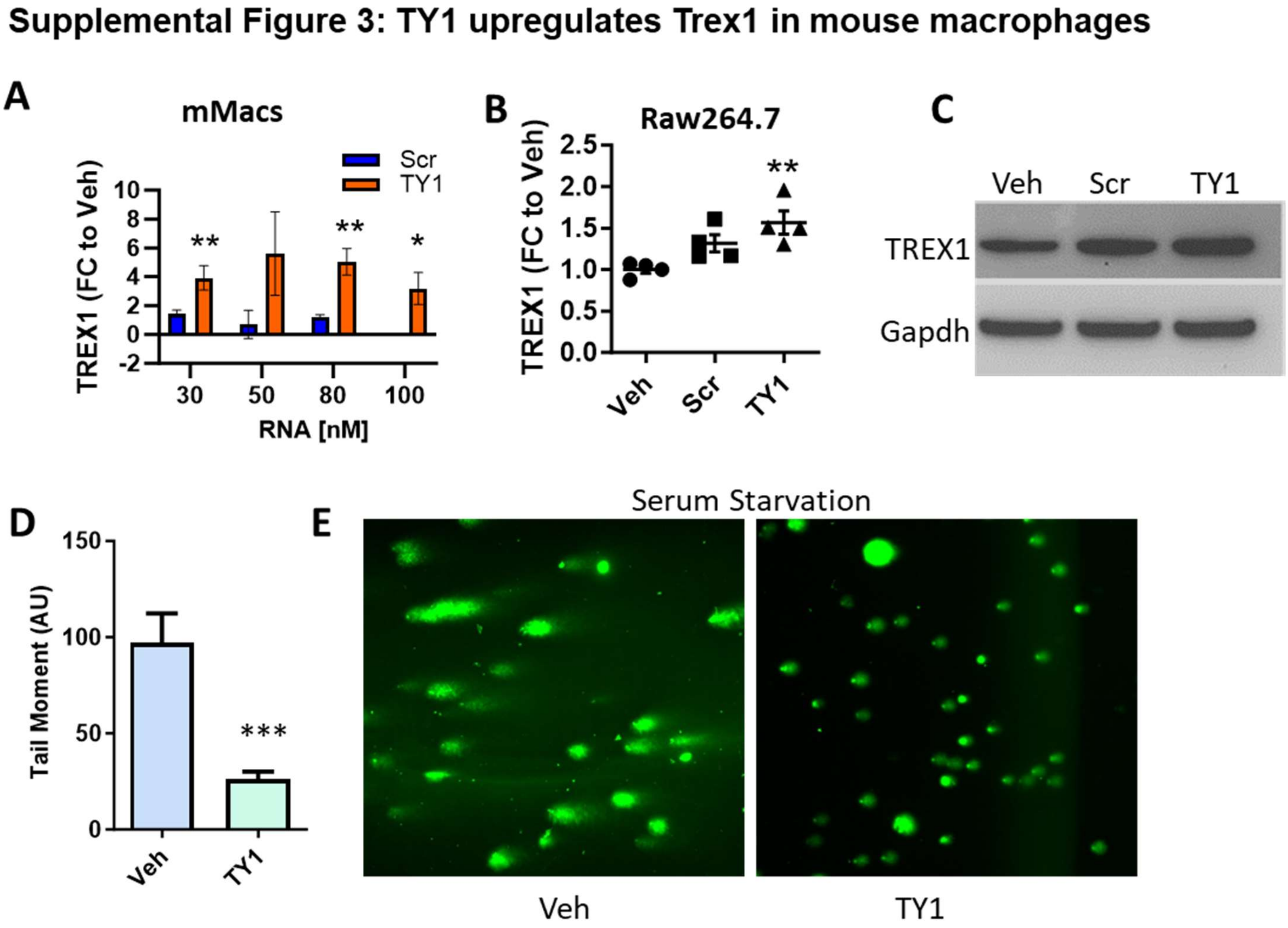


**Supplemental Figure 3: TY1 upregulates TREX1 in mouse macrophages.**
QPCR of mouse bone marrow-derived macrophages demonstrating dose-dependent upregulation of TREX1 with exposure to increasing concentration of TY1 (n=3 biological replicates per group). Analysis was done using Student's t test with 95% CI; *P<0.05; **, P<0.01; ***, P<0.001.

**Fig. S4.**

### Supplemental Figure 4: Chronic intravenous exposure of TY1 is well tolerated in healthy animals

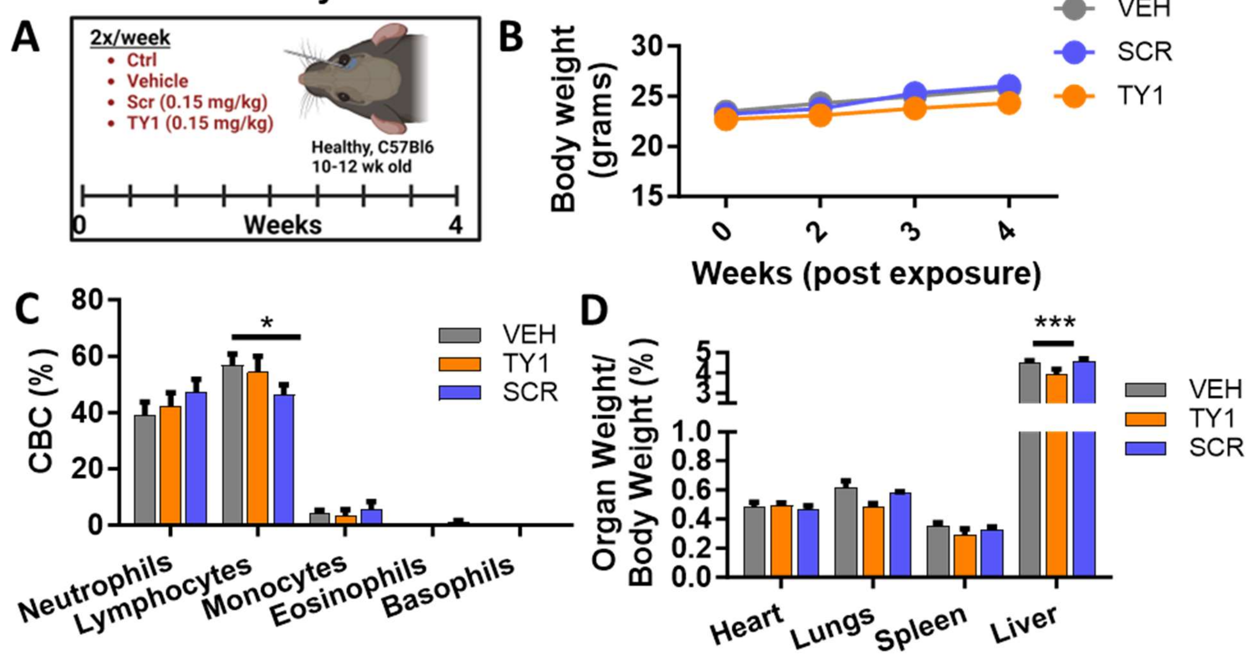


**E**

| Test | Control | Vehicle | Scramble | TY1 | Reference range |
|---|---|---|---|---|---|
| ALBUMIN | 1.9±0.2 | 1.9±0.1 | 1.9±0.4 | 2.1±0.1 | |
| AST (SGOT) | 54±29.7 | 45±7.9 | 34.7±4.6 | 48±20.8 | 10-45 IU/L |
| ALT (SGPT) | 25.3±15.4 | 31±15.3 | 16±3.6 | 28.3±24 | 10-35 IU/L |
| BUN | 18±3 | 17.3±1.2 | 18±3 | 18±3 | 9-30 mg/dL |
| Creatinine | 0.2±0.0 | 0.2±0.0 | 0.2±0.0 | 0.2±0.0 | 0.5-2.2 mg/dL |
| Glucose | 229.7±25.1 | 241.7±24.3 | 257.7±28 | 241.7±24.3 | 60-125 mg/dL (fasting) |
| Calcium | 6.5±0.5 | 6.8±0.0 | 6.6±1 | 6.9±0.4 | 8.0-12.0 mg/dL |
| Sodium | 149.3±2.1 | 150.3±1.2 | 152±4 | 149.7±1.2 | 140-160 mEq/L |
| Potassium | 4.3±1.3 | 4±0.4 | 4.5±0.8 | 5±1 | 4.3-5.8 mEq/L |
| Magnesium | 1.5±0.0 | 1.7±0.2 | 1.7±0.2 | 1.8±0.2 | |
| Triglycerides | 49±12.1 | 37.7±6.4 | 46.7±11.2 | 49.3±5 | |
| Cholesterol | 40.3±1.5 | 47±7.2 | 40.3±1.5 | 40.3±1.5 | 50-250 mg/dL |

**Supplemental Figure 4: Chronic intravenous exposure of TY1 is well tolerated in healthy animals.** (**A**) Study design for chronic intravenous (retro-orbital) exposure toxicity study (n=5 animals per group). (**B**) Animal weight profile over weeks of TY1 exposure. (**C**) Complete blood count, organ weights (**D**), and blood chemistry (**E**) of animals at four weeks. Bars represent group mean and error bars represent s.d. Significance was determined by one-way ANOVA; *P<0.05; **P<0.01; ***P<0.001. Scale bars: 100 µm.

**Fig. S5.**

### Supplemental Figure 5: TY1 increases TREX1 in macrophages in injured rat heart tissue.

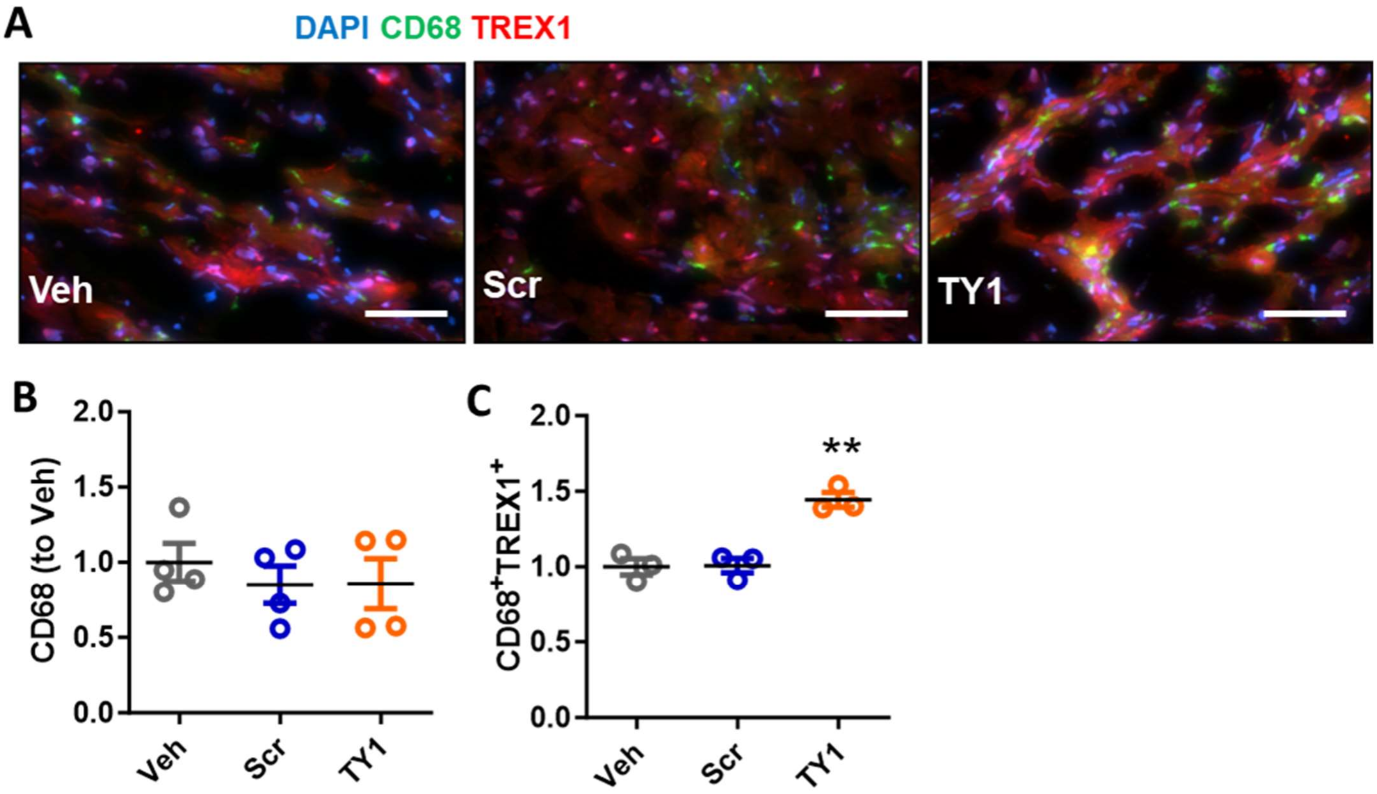


**Supplemental Figure 5: TY1 increases TREX1 in macrophages in injured rat heart tissue.** (**A**) Immunofluorescence of the left ventricles of MI animals (blue: DAPI, green: CD68 and TREX1: red, scale bar: 100 µm). Pooled data (n=3-4 animals per group) demonstrates comparable macrophage infiltration (CD68; **B**). TREX1 was also enriched in the macrophages in TY1-exposed animals compared to vehicle or scramble groups (**C**).

**Fig. S6**.

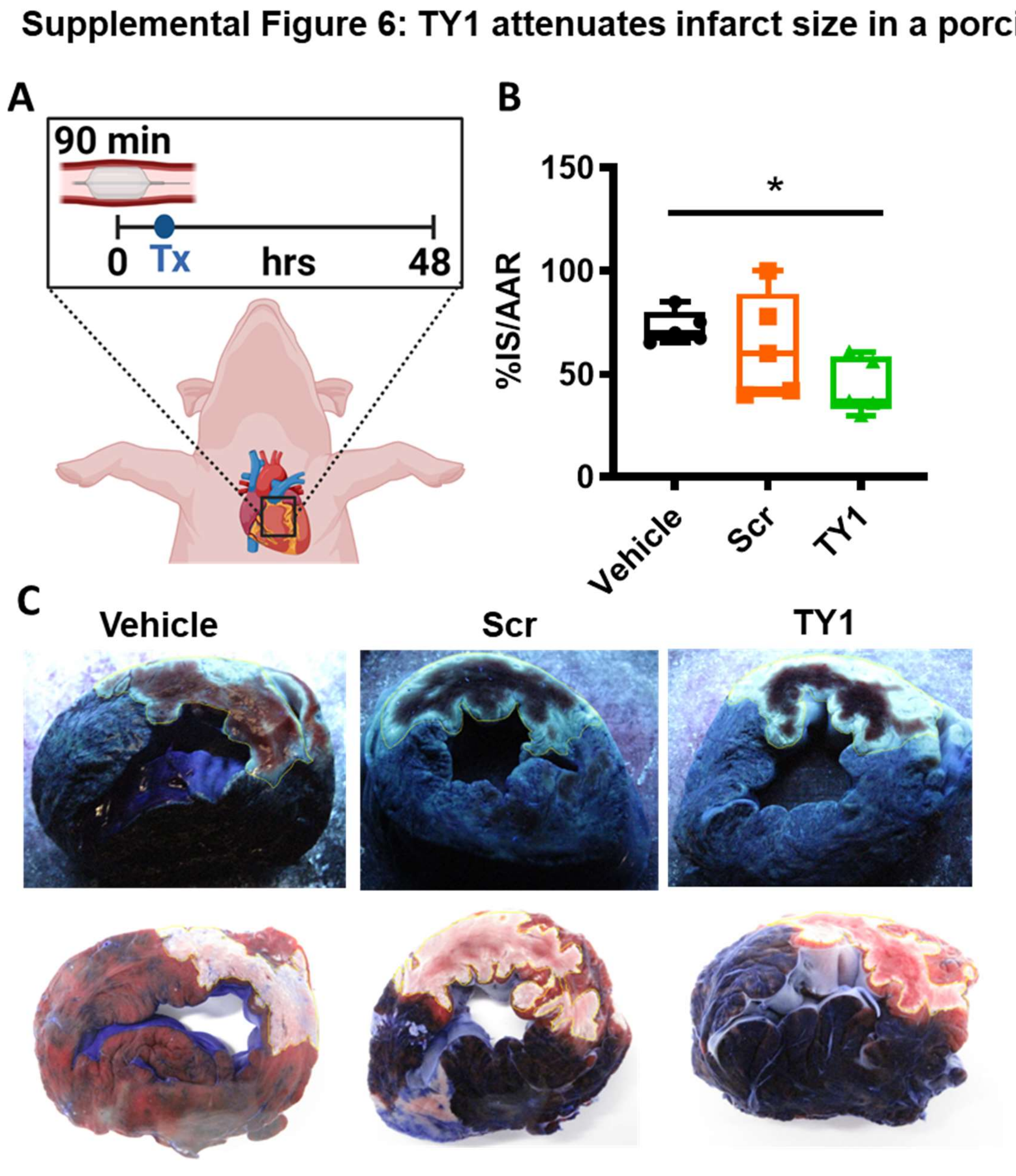


**Supplemental Figure 6: TY1 attenuates infarct size in a porcine model of MI**.
(**A**) Study design for TY1 administration in a porcine model of acute myocardial infarction (n=5 animals per group). Pigs received ischemia reperfusion injury (90 minutes of ischemia) followed by intravenous infusion of TY1, TY1 scrambled control (0.15 mg/kg), or vehicle. (**B, C**) Forty-eight hours post-injury, pigs receiving TY1 had reduced smaller infarct size to both scramble and vehicle. Lines represent group mean and error bars represent s.d. Significance was determined by one-way ANOVA; *P<0.05; **P<0.01; ***P<0.001.

**Table S1.**

**Supplementary Table 1: Primary assay IDs**

| Species | Gene | Assay ID |
| --- | --- | --- |
| Rat | Il10 | Rn01483988_g1 |
| Rat | IL6 | Rn01410330_m1 |
| Rat | P21 | Rn00589996_m1 |
| Rat | NF-kB | Rn01399572_m1 |
| Mouse | Il10 | Mm01288386_m1 |
| Mouse | P21 | Mm04205640_g1 |
| Human | Il10 | Hs00961622_m1 |
| Rat | TREX1 | Rn03062330_s1 |
| Mouse | TREX1 | Mm01229287_m1 |
